# Cardiac sympathetic denervation prevents sudden cardiac arrest and improves cardiac function by enhancing mitochondrial-antioxidant capacity

**DOI:** 10.1101/2023.01.29.526082

**Authors:** Swati Dey, Pooja Joshi, Brian O’Rourke, Shanea Estes, Deeptankar DeMazumder

## Abstract

**RATIONALE:** Sudden cardiac arrest (***SCA***) and heart failure (***HF***) are leading causes of death. The underlying mechanisms are incompletely understood, limiting the design of new therapies. Whereas most autonomic modulation therapies have not shown clear benefit in HF patients, growing evidence indicates cardiac sympathetic denervation (***CSD***) exerts cardioprotective effects. The underlying molecular and cellular mechanisms remain unexplored.

**OBJECTIVE:** Based on the hypothesis that mitochondrial reactive oxygen species (***mROS***) drive the pathogenesis of HF and SCA, we investigated whether CSD prevents SCA and HF by improving mitochondrial antioxidant capacity and redox balance, to correct impaired Ca^2+^ handling and repolarization reserve.

**METHODS AND RESULTS:** We interrogated CSD-specific responses in pressure-overload HF models with spontaneous SCA using *in vivo* echocardiographic and electrocardiographic studies and *in vitro* biochemical and functional studies including ratiometric measures of mROS, Ca^2+^ and sarcomere dynamics in left ventricular myocytes. Pressure-overloaded HF reduced mitochondrial antioxidant capacity and increased mROS, which impaired β-adrenergic signaling and caused SR Ca^2+^ leak, reducing SR Ca^2+^ and increasing diastolic Ca^2+^, impaired myofilament contraction and further increased the sympathetic stress response. CSD improved contractile function and mitigated mROS-mediated diastolic Ca^2+^ overload, dispersion of repolarization, triggered activity and SCA by upregulating mitochondrial antioxidant and NADPH-producing enzymes.

**CONCLUSIONS:** Our findings support a fundamental role of sympathetic stress-induced downregulation of mROS scavenging enzymes and RyR-leak mediated diastolic Ca^2+^ overload in HF and SCA pathogenesis that are mitigated by CSD. This first report on the molecular and cellular mechanisms of CSD supports its evaluation in additional high-risk patient groups.

**BRIEF SUMMARY:** Cardiac sympathetic denervation (***CSD***) confers cardioprotective effects via unclear mechanisms. In a guinea pig model that uniquely mimics human pressure-overload heart failure (***HF***) with spontaneous sudden cardiac arrest (***SCA***), we interrogated CSD-specific responses using echocardiographic, electrocardiographic and biochemical measures, and ratiometric measures of mitochondrial reactive oxygen species (***mROS***), Ca^2+^ and sarcomere dynamics. Consistent with our hypothesis, CSD rescued cardioprotection by upregulating mitochondrial antioxidant and NADPH-producing enzymes, which mitigate mROS-mediated Ca^2+^ derangements, repolarization lability, triggered activity, HF and SCA. Our findings provide the first molecular and cellular mechanistic basis for evaluating CSD therapy in a broader group of high-risk patients.

## INTRODUCTION

The sympathetic “fight-or-flight” stress response triggers the release of hormones that produce well-orchestrated changes throughout the body. Acute increases in sympathetic stress improve cardiovascular function. However, persistent sympathetic hyperactivity becomes maladaptive, increasing the risk of heart failure (***HF***), ventricular tachyarrhythmias (***VT***) and sudden cardiac arrest (***SCA***)(1). Despite optimal medical therapy, half of all HF patients die within the first five years of diagnosis and SCA accounts for about half of these deaths. The need for the design of new, more effective therapies for these two leading causes of death in the industrialized world remains a clinical, research and public health priority.

Autonomic dysfunction is a hallmark of both HF and SCA. However, pharmacological therapies aimed at autonomic targets have had limited success. Inhibitors of β-adrenergic signaling and the renin-angiotensin-aldosterone system (***RAAS***) delay mortality but do not reverse the disease process. Therapies for SCA are even more limited. While antiarrhythmic drugs may be effective at treating acute episodes of VT and SCA, they are proarrhythmic and increase mortality over the longer term. Moreover, β-adrenergic receptor blockers (***β-blockers***) do not reliably prevent SCA(2,3). As such, implantable cardioverter-defibrillators (***ICDs***) remain the only effective guideline-based therapy for primary and secondary prevention of SCA(4,5).

Progress in designing new therapies has been limited by poor insight into the underlying mechanisms. A growing body of evidence over the past century(6-8) indicates cardiac sympathetic denervation (***CSD***) protects against both HF(9-17) and SCA(17-21). The underlying molecular and cellular mechanisms are unknown, limiting CSD application to a small subset of high risk patients, i.e., those with long QT syndrome, catecholaminergic polymorphic VT, arrhythmogenic right ventricular cardiomyopathy, and VT episodes refractory to optimal medical therapy that require recurrent defibrillator shocks(8,12-16,19,22-25). An improved understanding of underlying CSD mechanisms could be used to increase its efficacy, extend its benefit to other high-risk patient groups and design new, more effective therapies targeting the pathways engaged by CSD.

We recently showed that mitochondrial reactive oxygen species (***mROS***) drive the pathogenesis of HF and SCA via distinct mechanistic pathways(26). Whether CSD engages these signaling pathways to confer salutary effects is unexplored. We hypothesized that CSD prevents SCA and HF by improving mitochondrial antioxidant capacity, autonomic and redox balance, Ca^2+^ handling and cardiac repolarization reserve. Our findings provide critical new insights into the role of mROS scavenging defects and diastolic Ca^2+^ in the pathogenesis and therapy of HF and SCA.

## METHODS

The foremost limitation for mechanistic studies of HF and SCA has been the lack of a suitable experimental model that mimics its human counterpart(27). Genetic models have contributed important mechanistic insight but most SCA victims do not have well-defined genetic abnormalities. Based on clinical guidelines, patients with poor left ventricular (***LV***) systolic function are eligible for primary prevention ICD implantation(4,5). However, the majority of SCA victims do not have advanced cardiovascular disease and remain asymptomatic until catastrophic clinical presentation. Non-ischemic etiologies for HF and VT are also more challenging to treat clinically because the underlying mechanisms are poorly understood. To address these unmet needs, we previously established a guinea pig pressure-overload model that mimics human HF and SCA(26). Unique to this model is a high incidence of spontaneous sustained VT that lead to SCA in freely ambulating animals. The VT episodes start occurring early during HF development. Transcriptomic, proteomic, metabolomic and functional studies demonstrated that this model recapitulates key features of human HF including derangements of autonomic, electrophysiological, hormonal, Ca^2+^ handling, redox, metabolic and inflammatory components as an integrative maladaptive response to cardiac insult. We further showed that high mROS levels cause chronic proteome remodeling, impairing contractility, and post-translational modifications of ion channels, triggering VT and SCA. Notably, the VT burden is highest after, but not during, mild transient β-adrenergic stress, as noted in humans(28-31). Thus, this model represents an ideal platform to investigate the salutary effects of CSD on HF and SCA. The *in vivo* (e.g., ECG, echocardiography), *in vitro* (e.g., biochemical and functional measures of mROS dynamics, mitochondrial antioxidant capacity, Ca^2+^ transients and sarcomere shortening) and statistical methods were described previously(26,32-34) and are detailed in the Supplementary Materials.

## RESULTS

We performed bilateral video-assisted thoracoscopic surgery to mimic the clinical CSD procedure for bilateral ablation and excision of the inferior half of the stellate ganglion (**Fig. 1A**), a discrete bundle of nerve cells near C7-T1 that provides the major sympathetic input to the heart. Surgical accuracy was confirmed by prototypical perioperative functional responses (**Fig. 1B**), i.e., graded changes in heart rate after sympathectomy (**Fig. 1-B**), and postoperative immunostaining for tyrosine hydroxylase (***TH***), neurofilament (***NF***) and 4’,6-diamidino-2-phenylindole (***DAPI***), which showed that the excised tissue section encompassed sympathetic neurons (**Fig. 1-C**).

**Figure 1.**
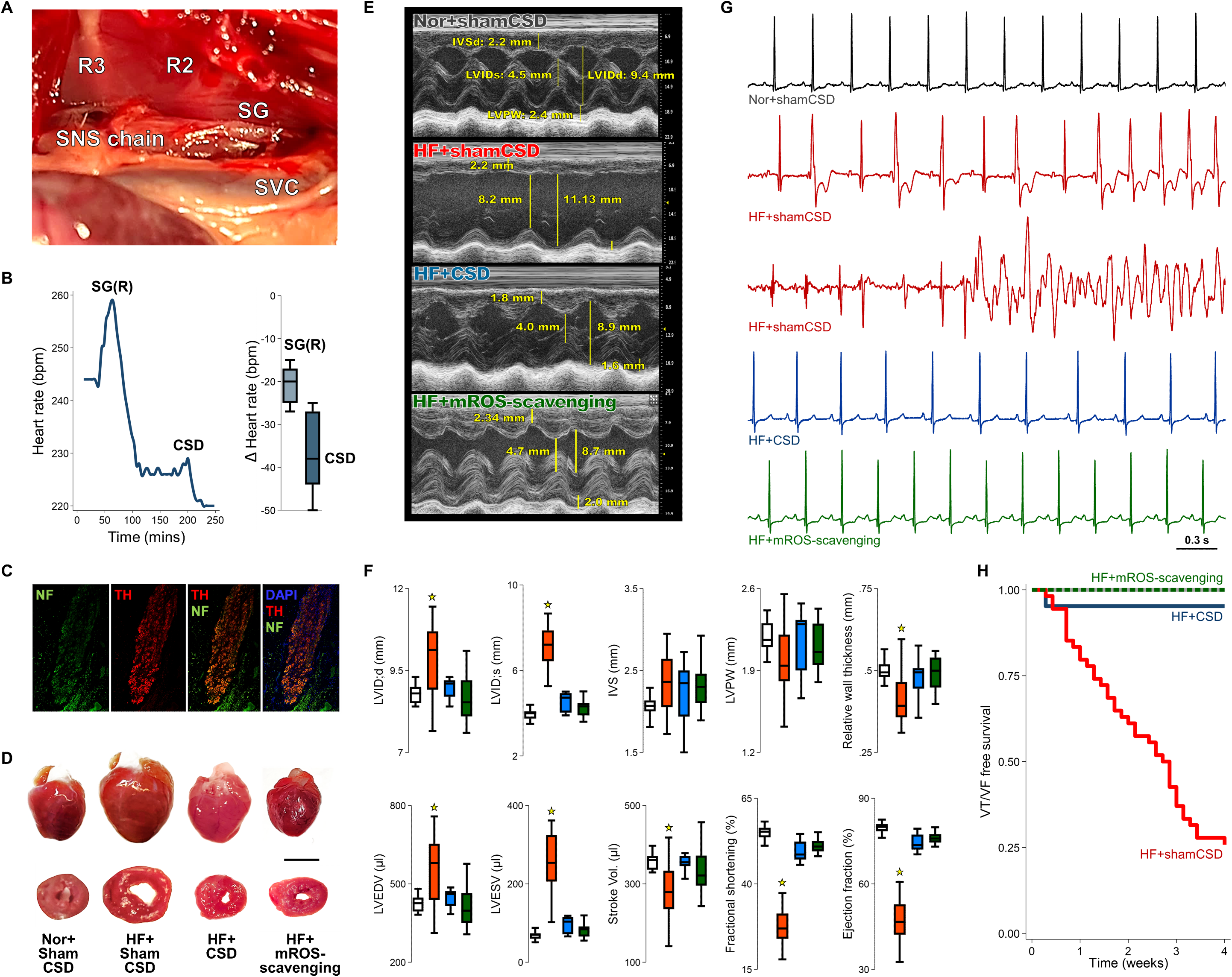
CSD prevents HF and SCA. **(A)** *In vivo* photo of guinea pig thoracic cavity showing the white fibrous sympathetic chain running along the anatomical landmarksrib 1 (R1), rib 2 (R2) and superior vena cava (SVC). The sSG, located between R2 and the spine, was accessed ventrally via the second intercostal space. **(B)** Representative example (left) of heart rate changes during right, SG(R), followed by left, SG(L), stellate ganglionectomy. The box plots (right) summarize steady-state heart rate changes. **(C)** Representative immunostaining for tyrosine hydroxylase (red) and neurofilament (green) with DAPI stained nuclei (blue) in excised SG tissue, confirming the presence of sympathetic neurons (bar=10 μm). **(D)** Representative gross heart (top) and corresponding cross-sections (bottom). **(E)** Representative M-mode echocardiography. **(F)** Echocardiographic systolic and diastolic LV internal dimensions (LVIDs) were larger in HF+shamCSD compared to HF+CSD and HF+mROS-scavenging animals. HF+shamCSD also had the lowest relative wall thickness (RWT), end-diastolic volume (EDV), end-systolic volume (ESV), fractional shortening (FS) and ejection fraction (EF) compared to HF+CSD and HF+mROS-scavenging animals, the FS and EF were greatest in normal+shamCSD animals. The ventricular septal (IVS) and LV posterior wall thickness (LVPW) were not different. **\***ANOVA-p<0.05; N≥15/group. **(G)** Representative ECGs showing spontaneous PVCs and VT in HF+shamCSD animals that were mitigated by CSD and mROS-scavenging. **(H)** The VT led to SCA in >70% of HF+shamCSD animals (N=68) that were mitigated by CSD (N=25) and mROS-scavenging (N=15) (two-tailed log-rank test).

The animals were randomized to five groups: (**1**) normal controls with sham CSD (***normal+shamCSD***); (**2**) normal controls with CSD (***normal+CSD***), (**3**) HF with sham CSD surgery (***HF+shamCSD***), (**4**) HF with CSD (***HF+CSD***) to abolish sympathetic hyperactivity, or (**5**) HF with continuous mROS scavenging *in vivo* with MitoTEMPO (***HF+mROS-scavenging***) for reducing global cellular oxidative stress(26,32). The primary endpoint was VT-free survival at 4 weeks. Because an *in vivo* model of HF+CSD with controlled increases in cardiac mROS does not currently exist, we interrogated LV myocytes from HF+shamCSD and HF+CSD models *in vitro* via acute exposure to isoproterenol for increasing endogenous ROS, and to graded amounts of exogenous ROS (i.e., H_2_O_2_) to measure antioxidant activity.

### CSD improves LV function

Serial echocardiography was performed in conscious animals to determine changes in cardiac structure and function (**Online Fig. 1**) and Doppler was used to confirm similar degrees of aortic banding over 4 weeks in HF+shamCSD and HF+CSD animals. In HF+shamCSD animals, aortic banding caused LV hypertrophy over the first 2 weeks that progressed to dilated cardiomyopathy over the next 2 weeks (**Fig. 1D-F**). By the end of 4 weeks, the animals exhibited features typical of human dilated cardiomyopathy such as increased LV systolic and diastolic volumes and decreased stroke volume, ejection fraction and fractional shortening. CSD improved upon these features, consistent with prior reports in humans(9-17). However, the ejection fraction and fractional shortening were still lower in HF+CSD compared to normal+shamCSD animals (**Fig. 1F**).

### CSD mitigates VT and SCA

Continuous wireless *in vivo* ECG analysis revealed a high burden of spontaneous premature ventricular contractions (***PVCs***) and VT in freely ambulating HF+shamCSD animals (**Fig. 1G**) that started as early as the first week, i.e., before onset of LV dysfunction, and led to SCA in >70% of animals by the fourth week (**Fig. 1H**). These findings are similar to humans in that most SCA victims do not have HF. As also noted in humans(17,19-21), CSD mitigated VT and SCA. These lifesaving effects of CSD were recapitulated by *in vivo* mROS scavenging (**Fig. 1E-H**), suggesting a common mechanistic pathway.

### CSD improves cardiac autonomic balance

In normal animals, CSD had no apparent effect on heart rate variability (***HRV***) and sympathovagal balance (**Online Fig. 2**), suggesting a large margin of safety for cardiac sympathetic input and dominant parasympathetic control of resting hearts. By contrast, hemodynamic compromise in HF+shamCSD animals increased heart rate and sympathetic tone, reduced HRV and total power, and blunted the chronotropic response to transient (1-hour) daily β-adrenergic challenge via an implanted automated wireless pump containing lowdose isoproterenol. Notably, VT and SCA episodes occurred more frequently over the hours following, but not during, β-adrenergic stress, as also seen in humans(28-31).

**Figure 2.**
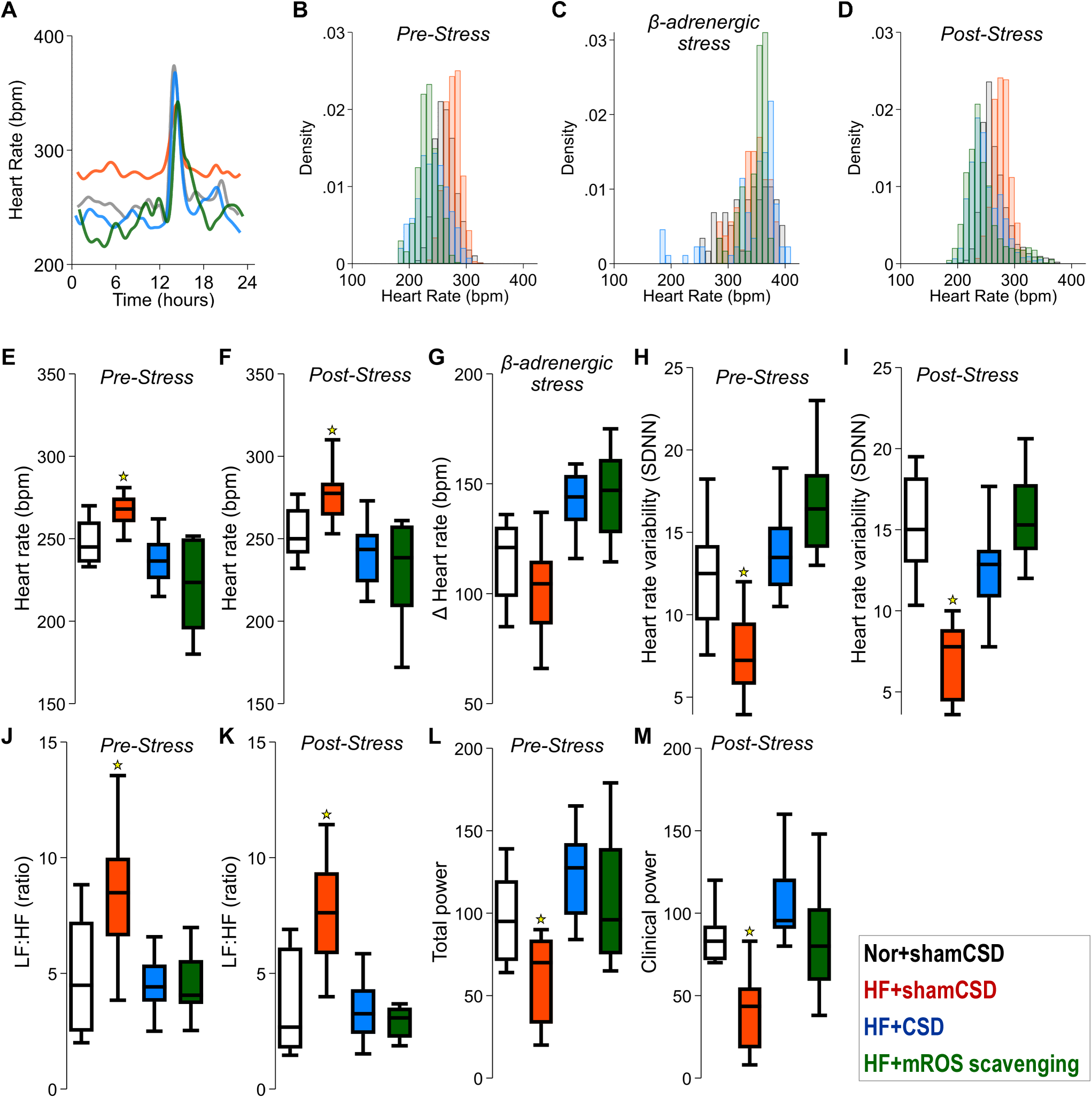
CSD improves autonomic balance and chronotropic competency. *p<0.01 vs. HF+shamCSD. Representative heart rates **(A)** and corresponding histograms **(B-D)** before (rest), during (stress), and 4 hours after (recovery) transient β-adrenergic stress with low-dose isoproterenol revealed that CSD (blue) and mROS-scavenging (green) lowered heart rates at rest and recovery and enhanced chronotropic response to stress (N≥7 animals/group). The box plots summarize the heart rates during rest (**E**) and recovery (**F**), chronotropic response during stress (**G**), heart rate variability during rest (**H**) and (**I**) recovery, the ratio of the low (LF) and high (HF) frequency bands of the heart rate power spectral density corresponding primarily to sympathetic and parasympathetic tone respectively during rest (**J**) and recovery (**K**), and total power for all frequency bands at rest (**L**) and recovery (**M**) (N>8 per group).

Despite pressure-overload induced hemodynamic compromise, CSD improved the heart rate (**Figs. 2A-G**), HRV (**Figs. 2H-I**), sympathovagal balance (LF/HF ratio; **Figs. 2J-K**), total power (**Figs. 2L-M**), and chronotropic response to β-adrenergic stress (**Fig. 2C,G**). Again, *in vivo* mROS scavenging yielded similar results to CSD (**Fig. 2**). Thus, we performed molecular and cellular studies to determine whether CSD reduces mROS levels and improves β-adrenergic signaling by enhancing mitochondrial antioxidant capacity.

### CSD improves redox balance by increasing mitochondrial antioxidant capacity

We tested this hypothesis by quantifying ROS dynamics in intracellular compartments of LV myocytes at the highest spatiotemporal resolutions(26,32). Briefly, we performed *in vivo* viral transduction of a genetically-encoded sensor, mitoORP, which is a ratiometric H_2_O_2_ sensor with dual-excitation wavelengths of 405/488 nm and emission at 520 nm. Each experiment was calibrated for the minimum and maximum wavelengths (405/488 nm ratio). The fractional oxidation of the sensor (F_ox_) was calculated to facilitate quantitative comparisons of basal and stimulated mROS levels across experimental groups.

The LV myocytes from pressure overloaded hearts exhibited high mROS levels (HF+shamCSD, F_ox_=0.59±0.02), which were mitigated by CSD (HF+CSD, F_ox_=0.38±0.03; p<0.01) (**Fig. 3A-C**). To determine whether CSD reduced mROS by increasing mitochondrial antioxidant capacity, we increased endogenous mROS generation by exposing the myocytes to isoproterenol (1 µM) with electrical field stimulation at 1 Hz. Compared to HF+shamCSD (F_ox_= 0.44±0.03), mROS levels remained much lower in HF+CSD myocytes (F_ox_=0.71±0.03; p<0.001) (**Fig. 3D**). We confirmed these findings by treating the myocytes with exogenous ROS (tert-butyl hydroperoxide, 1 mM). Again, mROS levels remained significantly lower in HF+CSD (F_ox_=0.48 ± 0.02) compared to HF+shamCSD myocytes (F_ox_=0.84 ± 0.03, p<0.001) (**Fig. 3E**). Compared to normal+shamCSD, LV myocytes from HF+CSD animals displayed slightly lower ROS scavenging capacities (p<0.01) consistent with their mildly impaired global LV function (**Fig. 1F**). These findings suggest that sympathetic hyperactivity is the primary, but not the only driver of the pathophysiology of pressure-overload HF and SCA.

**Figure 3.**
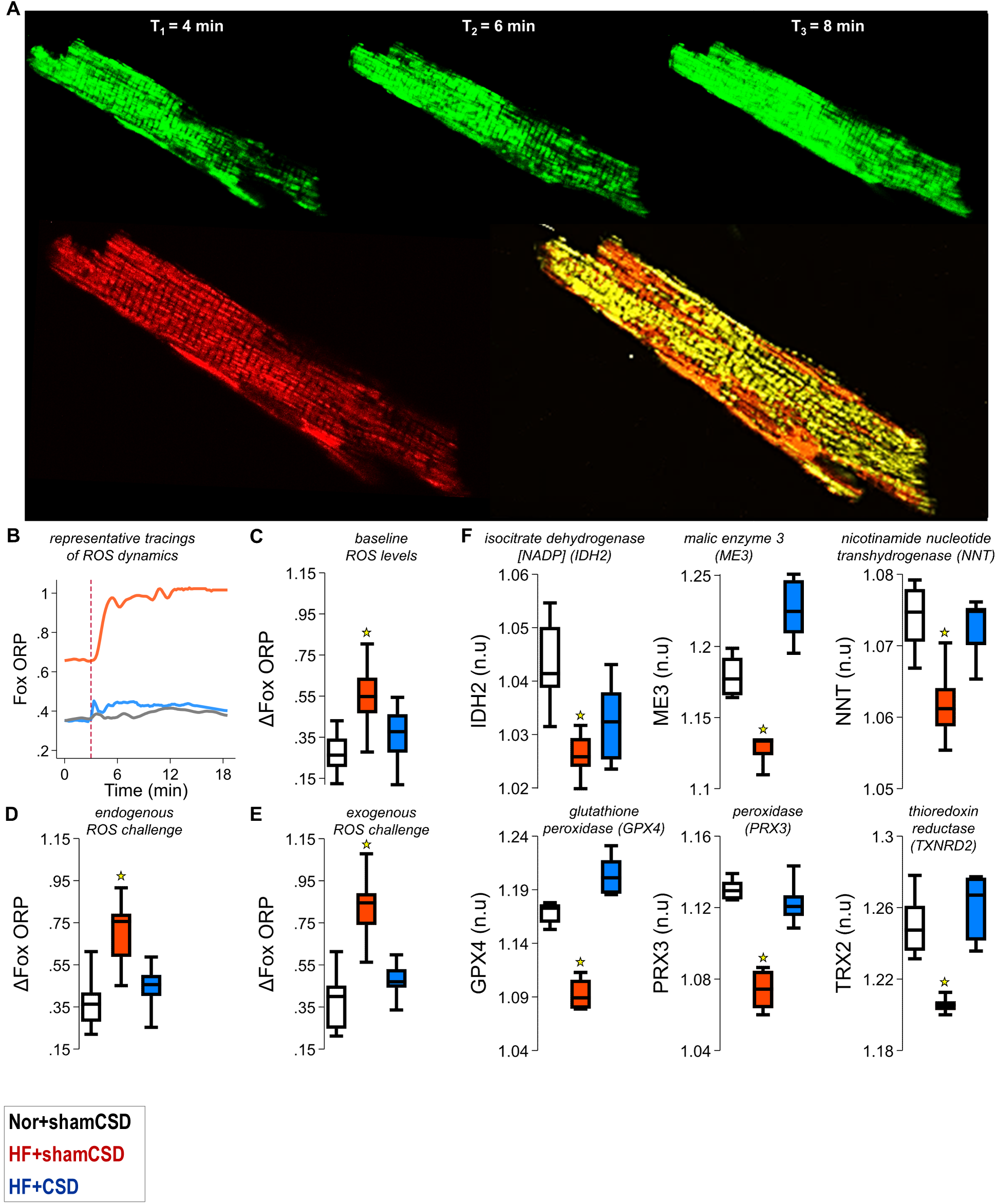
CSD enhances mitochondrial antioxidant capacity and reduces mROS levels in LV myocytes. **(A)** Representative images (top) of increasing GFP-mitoORP1 signal, indicating increasing ROS levels with isoproterenol during 1 Hz field stimulation. Co-localization (bottom) of mitoORP1 and TMRM indicates high specificity for mROS. **(B)** Corresponding time-lapse tracings of fractional oxidation (Fox) of GFP-mitoORP1 from normal+shamCSD (grey), HF+shamCSD (red) and HF+CSD (blue) are shown at baseline and after (vertically-dashed line) isoproterenol exposure. **(C)** Box plots show the highest baseline mROS in HF+shamCSD (p<0.001; N>10 myocytes/heart; >5 hearts/group). **(D)** Isoproterenol-induced endogenous mROS production (37°C; 1 Hz) increased baseline mROS by 10% in HF+CSD compared to 40% in HF+shamCSD myocytes despite already higher baseline values. Steady-state endogenous ROS were lower in HF+CSD than HF+shamCSD myocytes (p<0.001). **(E)** Exogenous oxidative stress by 1mM tert-butyl hydroperoxide revealed impaired scavenging in HF+shamCSD compared to HF+CSD myocytes, indicating CSD improves mitochondrial antioxidant capacity (p<0.001). The HF+CSD group displayed a small but significantly lower ROS scavenging capacity compared to normal+shamCSD (p<0.01). **(F)** qPCR results show that mitochondrial antioxidant enzymes in the thioredoxin and glutathione antioxidant pathways were decreased by pressure overloaded HF and CSD upregulated these enzymes (N>5 hearts/group).

To identify the molecular basis for CSD-mediated preservation of mitochondrial antioxidant capacity, we quantified mRNA expression of mitochondrial antioxidant and NADPH-producing enzymes (**Fig. 3F**). CSD increased the expression of isocitrate dehydrogenase [NADP] (***IDH2***), glutathione peroxidase (***GPX4***), peroxidase (***PRX3***), thioredoxin reductase (***TXNRD2***), malic enzyme 3 (***ME3***) and nicotinamide nucleotide transhydrogenase (***NNT***) that otherwise were reduced by pressure-overload HF. These findings indicate that reduced mROS scavenging is a critical pathogenic component of HF, and that CSD exerts its cardioprotective effects, at least in part, by enhancing mitochondrial antioxidant capacity and reducing mROS levels in LV myocytes. We previously showed that high mROS levels control global cellular oxidative stress, leading to widespread damage including derangements in Ca^2+^ handling and β-adrenergic receptor-mediated signaling(26,32). Further, ROS oxidizes thiol groups of cysteine residues on ryanodine receptors (***RyRs***)(35-37), augmenting spontaneous Ca^2+^ leak from the sarcoplasmic reticulum (***SR***) during diastole(37) that in turn, increases diastolic Ca^2+^ levels(38). High diastolic Ca^2+^ activates a transient inward current involving primarily the Na-Ca exchanger, resulting in delayed afterdepolarizations (***DADs***), which can induce additional triggered activity (i.e., early afterdepolarizations) and repolarization lability, leading to VT and SCA. Indeed, the arrhythmogenic effects of cardiac glycosides involve mROS generation and RyR oxidation(39). Whether ROS induced by sympathetic hyperactivity and its downstream effects on SR Ca^2+^ handling can be reversed by CSD has not been previously studied.

### CSD improves SR Ca^2+^ load, β-adrenergic responsiveness and sarcomere shortening

The HF+shamCSD LV myocytes exhibited Ca^2+^ transients with slower upstrokes (reflecting reduced SR Ca^2+^ load), blunted peaks, prolonged decays (**Fig. 4A-E**), and blunted responses to maximal β-adrenergic receptor activation (**Fig. 4F-I**), along with similar impairments in sarcomere shortening (**Fig. 4J-R**). CSD improved Ca^2+^ transient upstroke, peaks and decays. The consequent faster and larger Ca^2+^ transients, in the absence and presence of β-adrenergic stimulation, improved myofilament contraction, as evidenced by increased sarcomere shortening (**Fig. 4J-R**) and accounting for the increase in global LV systolic function (**Fig. 1E-F**).

**Figure 4.**
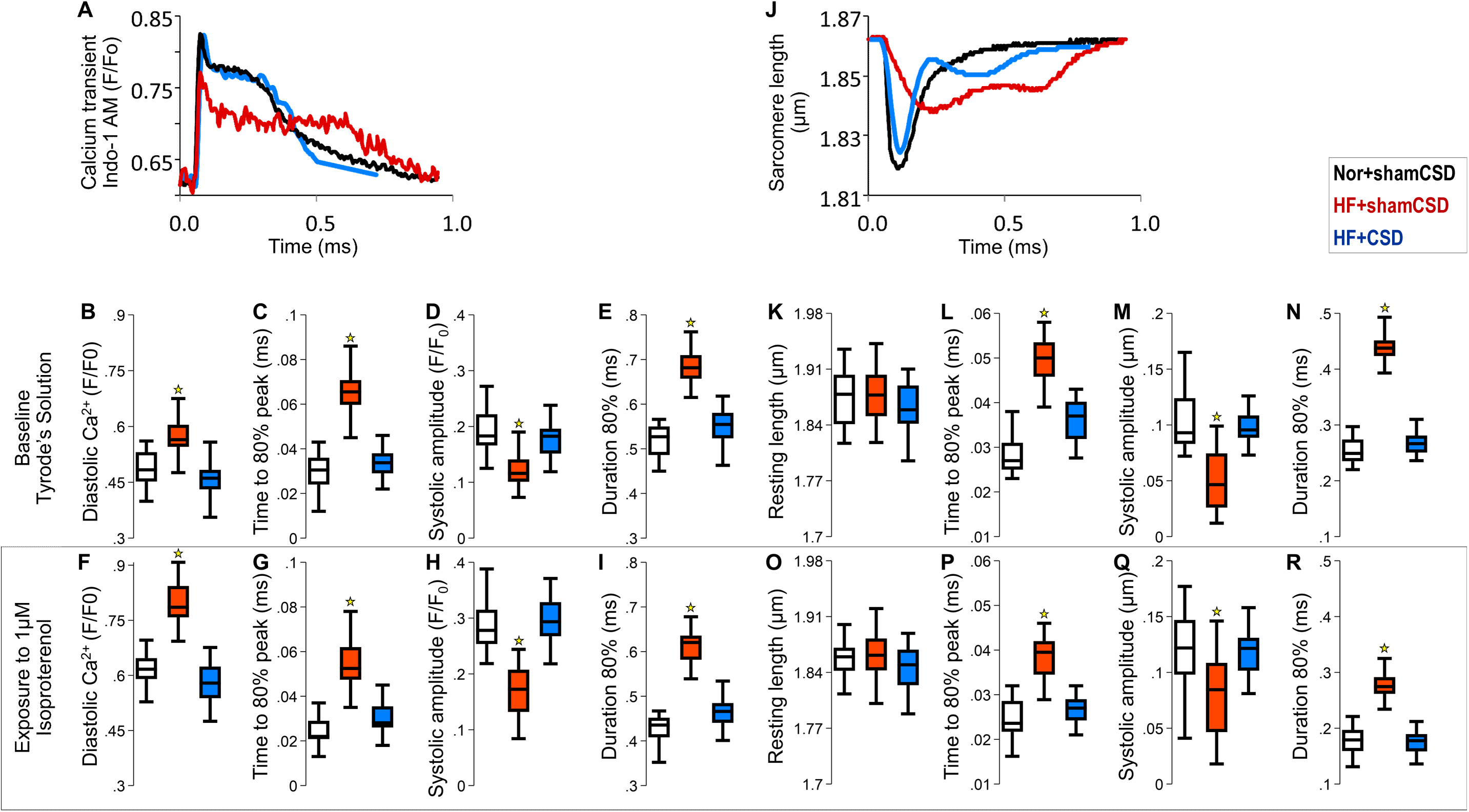
CSD improves Ca^2+^ handling, sarcomere shortening and β-adrenergic responsiveness in LV myocytes. Representative Ca^2+^ transients normalized to resting diastolic values **(A)** and their derived features are shown before **(A-E)** and after **(F-I)** 1 μM isoproterenol exposure (37°C; 1 Hz). The small secondary increases during relaxation, sometimes seen due to L2 mode of Ca^2+^ channel gating(33), did not affect the results. Corresponding sarcomere shortening results are shown **(J-R)**. (N>10 myocytes/heart; >5 hearts/group; **\***p<0.01 vs. HF+shamCSD).

### CSD mitigates ROS-induced SR Ca^2+^ leak, diastolic Ca^2+^ overload, dispersion of Ca^2+^ transients, dispersion of repolarization, triggered activity, VT and SCA

To determine the mechanistic link between ROS, SR Ca^2+^ leak, diastolic Ca^2+^, and arrhythmic risk, we induced endogenous ROS generation or added exogenous ROS to LV myocytes and quantified the incidence of disturbances in Ca^2+^ transients and sarcomere shortening, such as delayed after-transients and after-contractions (**Fig. 5A**). Further, we quantified beat-to-beat dispersion in Ca^2+^ transient duration (***dCaTd***) by its coefficient of variation, calculated as 5-transient bins of the mean divided by the standard deviation of the time from the Ca^2+^ transient peak to the time to return to 80% diastolic Ca^2+^ levels (**Fig. 5B**). These disturbances occurred more frequently in HF+shamCSD compared to HF+CSD myocytes (25% vs. 5%; p<0.001) and were reversibly suppressed by *in vitro* ROS scavenging (**Fig. 5B-C**). Whereas increasing ROS increased diastolic Ca^2+^, scavenging ROS lowered pacing-induced diastolic Ca^2+^ (**Fig. 5B-C**), suggesting a direct mechanistic link. Mitochondrial ROS was the upstream effector of RyR-mediated leak and increases in diastolic Ca^2+^ because when caffeine increased RyR leak **(Fig. 5-B)**, mROS scavenging was no longer effective. Dantrolene, which stabilizes RyRs to prevent SR Ca^2+^ leak during diastole, eliminated these disturbances despite high mROS levels, indicating that SR Ca^2+^ leak via RyRs mediated the triggered activity.

**Figure 5.**
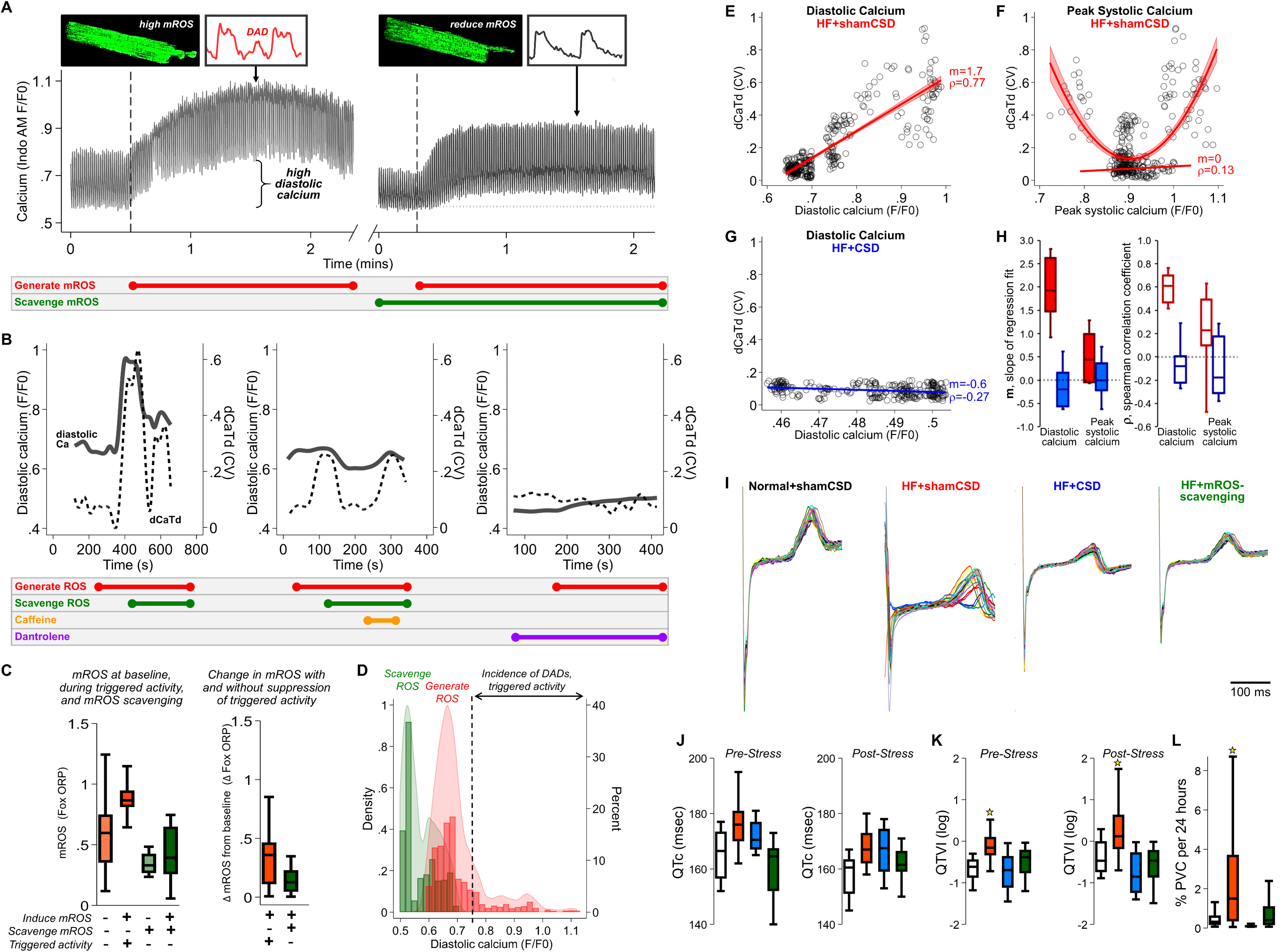
CSD reduces triggered activity and dispersion of repolarization by reducing ROS-induced SR Ca^2+^ leak-mediated diastolic Ca^2+^ overload. Snapshots of GFP-mitoORP1 signal (top) are shown in a HF+shamCSD myocyte during increases in diastolic Ca^2+^ and dCaTd that were followed by DADs and triggered activity within 1 minute of ROS generation by pacing and isoproterenol exposure (left panel); these effects were prevented by mitoTEMPO (right). **(B)** ROS scavengers also reduced diastolic Ca^2+^ and dCaTd (left), demonstrating a direct relationship. Caffeine, which increases RyR open probability, reversibly increased diastolic Ca^2+^ and dCaTd despite the presence of ROS scavengers (middle). Dantrolene, which stabilizes RyRs during diastole, reduced diastolic Ca^2+^ and dCaTd despite increasing ROS levels (right). **(C)** Summary results from HF+shamCSD myocytes show that mROS increased by about 30% from baseline (orange) before onset of triggered activity (red). ROS scavenger (green) abolished the high mROS levels, abnormal Ca^2+^ levels, dCaTD and triggered activity (N=20 myocytes from 5 hearts). Similar experiments could not be performed in normal, HF+CSD and HF+mROS-scavenging myocytes because they had lower mROS levels and rarely exhibited triggered activity. **(D)** The distributions of diastolic Ca^2+^ levels from all cells during ROS generation and scavenging revealed that triggered activity occurred frequently when diastolic Ca^2+^ exceeded a threshold (dashed line). **(E)** Representative example of the strong association between diastolic Ca^2+^ levels and dCaTd in HF+shamCSD myocytes despite the temporal lag in dCatTd changes with increasing diastolic Ca^2+^ (Spearman correlation coefficient, ***ρ***=0.77; slope of linear regression fit, ***m***=1.7). **(F)** Representative example of weakly positive correlation between peak systolic Ca^2+^ and dCaTd in HF+shamCSD myocytes, likely from impaired mitochondrial calcium loading and consistent with a better polynomial fit. **(G)** Representative example of weakly inverse correlation between peak diastolic Ca^2+^ and dCaTd in HF+CSD myocytes. **(H)** The corresponding summary of ***ρ*** and ***m*** values are shown for HF+shamCSD (red; N=90 myocytes from 5 hearts) and HF+CSD (N=75 myocytes from 5 hearts). **(I)** ECG R-to-T segments over 24 hours show increased dispersion of repolarization in HF+shamCSD that were reduced by CSD and mROS-scavenging. **(J)** The corrected QT intervals (QTc) were similar between HF+shamCSD and HF+CSD at rest and during recovery from stress, but prolonged compared to normal+shamCSD (p<0.01). **(K)** CSD reduced the high QT variability at rest and recovery of HF+shamCSD animals (n=>8 animals/group, p<0.01). **(L)** CSD also lowered the 24-hour PVC burden of HF+shamCSD animals.

In HF+shamCSD myocytes, the high mROS levels strongly correlated with pacing-induced diastolic Ca^2+^ levels (**Fig. 5D**), which strongly correlated with dCaTd (**Fig. 5E**) and triggered activity when pacing-induced diastolic Ca^2+^ and dCaTd were high. By contrast, dCaTd and triggered activity correlated poorly with peak systolic Ca^2+^ (**Fig. 5F**). These findings suggest diastolic Ca^2+^ drives dCaTd and triggered activity. CSD abolished these associations and markedly lowered mROS (**Fig. 3A-C**), pacing-induced diastolic Ca^2+^ and dCaTd (**Fig. 5G-H**).

Consistent with the *in vitro* findings, HF+shamCSD animals exhibited prolonged repolarization and increased dispersion of repolarization as indexed by beat-to-beat changes in the ECG’s QT interval (**Fig. 5I**), in association with higher VT incidence (**Fig. 1G-H**). Notably, CSD did not shorten repolarization (**Fig. 5J**) but reduced its dispersion (**Fig. 5K**) and reduced 24-hour PVC burden (**Fig. 5L**). In a prior multicenter prospective study of HF patients, we showed that fluctuation of repolarization strongly predicts SCA, independent of other clinical parameters including QT prolongation(34).

## DISCUSSION

Despite growing clinical use(6-24), the molecular and cellular mechanisms underlying the benefits of CSD therapy remain unexplored. A key mechanistic finding in the present study is that CSD enhances mitochondrial antioxidant capacity by upregulating mitochondrial antioxidant and NADPH-producing enzymes in the setting of pressure-overload HF. The consequent increase in mROS scavenging reduces global oxidative stress, reducing RyR-mediated SR Ca^2+^ leak that in turn, improves sarcoplasmic Ca^2+^ load, β-adrenergic responsiveness and global LV function. Another key finding is that the high mROS-induced diastolic Ca^2+^ drives the beat-to-beat variability of Ca^2+^ transients and triggered activity. Diastolic Ca^2+^ levels are more important than sarcoplasmic Ca^2+^ load for driving the beat-to-beat variability of Ca^2+^ transients and triggering arrhythmogenic activity. The instability of diastolic Ca^2+^ handling may further exacerbate mROS in a vicious cycle(40). This is the first report that CSD mitigates VT and SCA by reducing mROS-induced diastolic Ca^2+^-mediated beat-to-beat Ca^2+^ transient variability, trigged activity and dispersion of repolarization (**Fig. 6 – Central Illustration**). Our findings support the clinical evaluation of CSD in a broader group of high-risk patients such as those with non-ischemic HF, early stages of LV dysfunction or high SCA risk who are not eligible for primary prevention ICDs.

**Figure 6.**
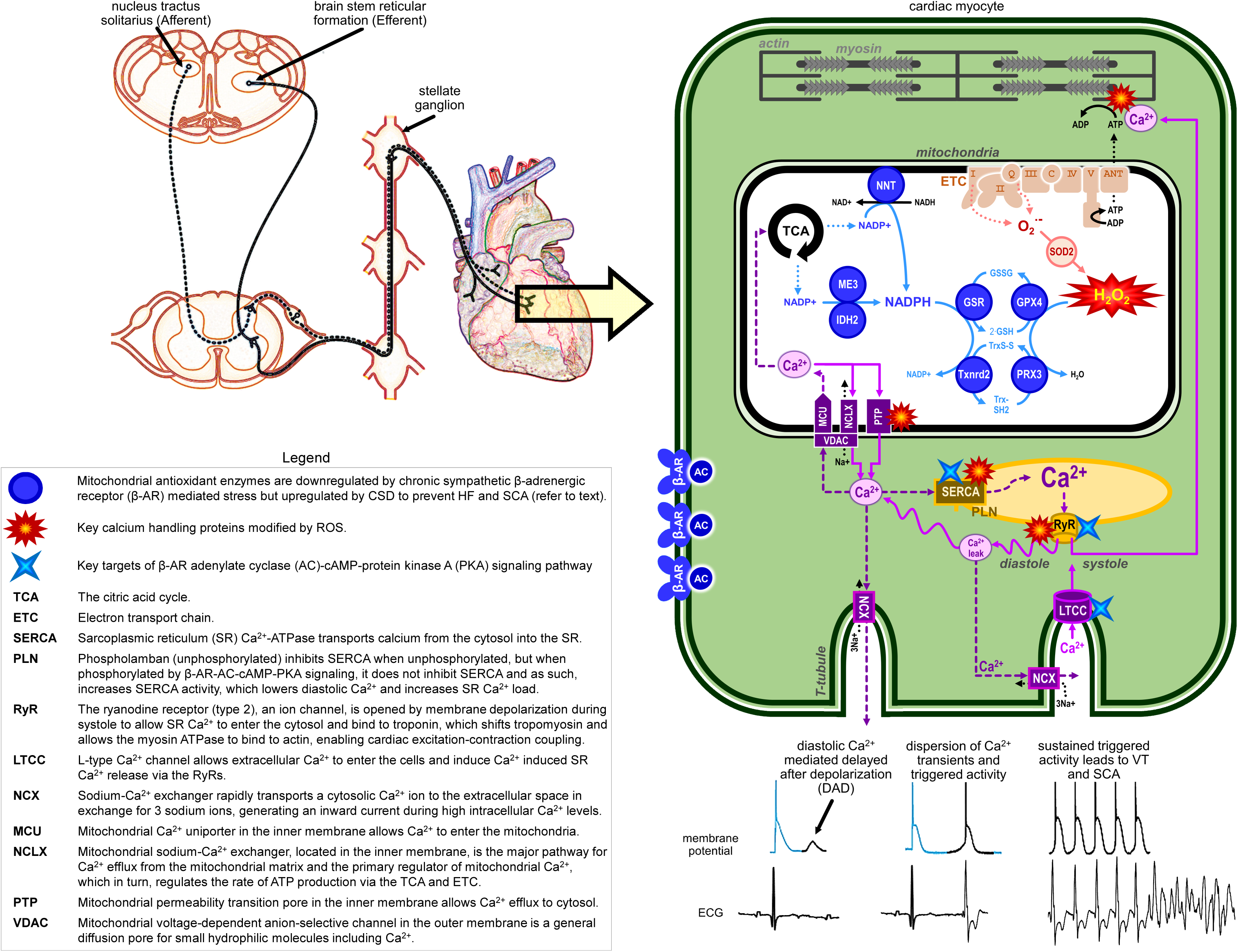
CENTRAL ILLUSTRATION: Chronic sympathetic stress-induced downregulation of mitochondrial antioxidant enzymes causes mROS-induced SR Ca^2+^ leak from RyRs and diastolic Ca^2+^ overload that drive SCD and HF pathogenesis; CSD targets these processes to rescue cardioprotection.

Autonomic dysfunction is a hallmark of HF and SCA(1,17,19-21,33). Although β-blockers delay HF-associated mortality, they do not reverse the underlying disease. Moreover, β-blockers do not reverse functional status in HF patients(28). β-blockers are also inadequate for primary and secondary prevention of SCA(2,3). Thus, ICDs remain the first-line guideline-recommended therapy for primary and secondary prevention of SCA in HF patients receiving optimal medical therapy with β-blockers, SGLT2 inhibitors and RAAS antagonists(4,5). In fact, initiation or escalation of β-blocker therapy for terminating VT is not recommended in HF patients because acutely, β-blockers can worsen HF. By contrast, CSD prevents VT refractory to β-blockers in HF patients. Thus, the goal of the present study was to interrogate the effects of CSD on pressure-overloaded hearts without the confounding effects of less effective therapies for VT.

During acute stress, sympathetic nerves originating from the SG release norepinephrine to stimulate cardiac β-adrenergic receptors on LV myocytes, activating adenylate cyclase–cAMP– protein kinase A signaling, which phosphorylates excitation-contraction coupling proteins such as SERCA, phospholamban, L-type Ca^2+^ channels and RyRs. The consequent increase in Ca^2+^ flux and SR Ca^2+^ load, along with synchronization of SR Ca^2+^ release during systole, elicits positive inotropy to meet physiological demands. By contrast, persistent activation of β-adrenergic receptors becomes maladaptive(26), increasing energy demand and workload that in turn, generate mROS, a primary source of oxidative stress in the heart(26). The consequent increase in ATP and NADPH consumption weakens mitochondrial antioxidant defense.

The findings of the present study show that mROS scavenging capacity is further reduced by downregulation of mitochondrial antioxidant enzymes. Beyond a certain threshold, as the rate of mROS generation exceeds the rate of mROS scavenging, the excess mROS released from a mitochondrion propagate to and induce ROS generation in other mitochondria and intracellular compartments. This regenerative positive-feedback cycle of ROS-induced ROS release increases global cellular ROS and oxidizes multiple ion channels and transporters. The RyRs, in close proximity to, and tightly coupled with, the mitochondrial network, are readily modified by ROS(35-37) via direct oxidation of thiol groups of cysteine residues(35,37) and Ca^2+^-independent activation of CaMKII, that increase SR Ca^2+^ leak. The consequent increase in diastolic Ca^2+^ causes dispersion of Ca^2+^ transients, dispersion of membrane repolarization and triggered activity, promoting VT and SCA. The spatiotemporal dispersion of repolarization between regions with the highest Ca^2+^ load (generating spontaneous SR Ca^2+^ release) and lower Ca^2+^ load may also promote VT. The depletion of SR Ca^2+^ load also blunts the rapid rise in Ca^2+^ in the diadic space, impairing mitochondrial Ca^2+^ signaling and increasing mROS.

Our findings demonstrate that CSD mitigates this vicious ROS-mediated cycle of widespread damage and protects against HF and SCA by: 1) increasing mROS scavenging rate by upregulating mitochondrial antioxidant and NADPH-producing enzymes; 2) reducing mROS-induced SR Ca^2+^ leak, thereby improving SR Ca^2+^ load, β-adrenergic signaling and chronotropic competency, and improving LV contractility; and 3) mitigating diastolic Ca^2+^ overload, dispersion of Ca^2+^ transients and repolarization, and SCA risk.

Although our findings also show that scavenging mROS *in vivo*, another potential therapeutic strategy(41,42), confers salutary effects similar to those of CSD, it is noteworthy that ROS are not only involved in pathological processes but ROS also participate in normal physiological signaling(43). Ca^2+^ handling in healthy cardiomyocytes is especially sensitive to ROS levels, such as RyR activation, SERCA inhibition and modulation of L-type Ca^2+^ channel function. Even small increases in mROS have important physiological roles such as increasing mitochondrial biogenesis, regulating mitochondrial respiration, lipolysis and differentiation, and inducing proton leak without damaging the mitochondrial membrane. As such, excessive ROS scavenging may prevent SCA but can also disturb signaling processes that are beneficial for cell function. For example, excessive overexpression of mitochondrial catalase is linked to impaired mitophagy(44).

The present findings indicate that CSD lowers oxidative stress without limiting the necessary ROS-dependent physiological processes. The CSD-mediated engagement of these targets provides strong mechanistic basis to further evaluate its clinical efficacy in other high-risk populations. Further, modulating sympathovagal tone via implantable nerve stimulators for modulating cardiac oxidative stress and its downstream effects on cardiac transcriptome, proteome and post-translational remodeling could be a viable, safe and transient/reversible therapeutic intervention, aimed at improving contractile function, preventing life-threatening arrhythmias and opening new avenues for enhancing human health.

### Limitations

Surgical CSD was performed at the onset of pressure-overloaded HF. High risk patients are typically brought to clinical attention after HF onset. We previously demonstrated(26) in a cross-over study of HF animals treated with *in vivo* mitoTEMPO that scavenging mROS *reverses* both HF and SCA risk. CSD likely engages the same mechanistic pathway after the onset of HF as that reported herein at the onset of HF. Furthermore, the risk of SCA is especially high during early stages of LV remodeling, i.e., before LV function has declined. The majority of SCA victims have little or no impairment in LV function and thus, are not eligible for primary prevention ICD implantation. These patients are also more challenging to study because they are routinely misclassified as low risk by current clinical criteria(4,5). The findings of the present study provide important mechanistic insight into this larger and underexplored high-risk group without HF.

## Acknowledgements

SD and DD jointly conceived and designed this study and wrote the manuscript. All authors reviewed and revised the manuscript.

## ABBREVIATIONS

CSD: Cardiac sympathetic denervation
Cyto-ORP: Cytosolic H_2_O_2_ probes
DAD: Delayed afterdepolarizations
DAPI: 4’,6-diamidino-2-phenylindole
dCaTd: Beat-to-beat fluctuations in the relaxation phase of Ca^2+^ transients
HF: Heart failure due to pressure-overload from ascending aortic banding
HF+CSD: Animal group subjected to HF and CSD
HF+mROS-scavenging: Animal group subjected to HF and continuous MitoTempo treatment
HF+shamCSD: Animal group subjected to HF and Sham surgery for CSD
HRV: Heart rate variability
LTCC: L-type voltage-dependent calcium channel
LV: Left ventricular
Mito-ORP: Mitochondrial ratiometric probes for measuring H_2_O_2_ levels
mROS: Mitochondrial reactive oxygen species
normal+CSD: Normal control animal group subjected to CSD
normal+shamCSD: Normal control animal group subjected to Sham surgery for CSD
ORP1: Yeast peroxidase
RAAS: Renin-angiotensin-aldosterone system
PVC: Premature ventricular contraction
RyR: Ryanodine receptors
SCA: Sudden cardiac arrest
SR: Sarcoplasmic reticulum
VT: Ventricular tachyarrhythmias

## CLINICAL PERSPECTIVE

### Competency in Medical Knowledge

Free radicals originating from the mitochondria of heart cells can cause widespread damage and drive the pathophysiology of heart failure, ventricular tachyarrhythmias and sudden cardiac arrest, the leading causes of death in the industrialized world. Targeted surgical excision of stellate ganglia improves cardiac systolic and diastolic function and protects against ventricular tachyarrhythmias and sudden cardiac arrest by enhancing mitochondrial antioxidant capacity and reducing free radicals throughout the heart.

### Translational Outlook 1

Because high oxidative stress is central to the pathophysiology of heart disease, modification of stellate ganglia activity has potential for broader application, i.e., beyond the limited subset of patients currently eligible for cardiac sympathetic denervation.

### Translational Outlook 2

Further clinical studies are needed to investigate whether targeted stellate ganglia modification will favorably impact outcomes in patients with different types of cardiomyopathies, arrhythmias and heart disease.

### Twitter

Free radicals originating from the mitochondria of heart cells can cause widespread damage and drive the pathophysiology of heart failure and sudden cardiac arrest, the leading causes of death in the industrialized world. Cardiac sympathetic denervation enhances mitochondrial antioxidant capacity and reduces free radicals throughout the heart.

## REFERENCES

1. DeMazumder D, Tomaselli GF. Molecular and cellular mechanisms of cardiac arrhythmias. In: Hill JA, Olson, E.N., editor Muscle 2-Volume Set: Fundamental Biology and Mechanisms of Disease. Philadelphia, P.A.: Elsevier Inc., 2012.

2. Arnold SV, Silverman DN, Gosch K et al. Beta-Blocker Use and Heart Failure Outcomes in Mildly Reduced and Preserved Ejection Fraction. JACC: Heart Failure 2023.

3. Al-Gobari M, El Khatib C, Pillon F, Gueyffier F. beta-Blockers for the prevention of sudden cardiac death in heart failure patients: a meta-analysis of randomized controlled trials. BMC Cardiovasc Disord 2013;13:52.

4. Gehi A, Haas D, Fuster V. Primary prophylaxis with the implantable cardioverter-defibrillator: the need for improved risk stratification. JAMA 2005;294:958–60.

5. Bardy GH, Lee KL, Mark DB et al. Amiodarone or an implantable cardioverter-defibrillator for congestive heart failure. N Engl J Med 2005;352:225–37.

6. Leriche R, Herman L, Fontaine R. Ligature de la coronaire gauche et fonction du coeur après énervation sympathique. CR Soc Biol 1931;107:547.

7. Cox W, Lewiston M, Robertson H. The effect of stellate ganglionectomy on the cardiac function of intact dogs: and its effect on the extent of myocardial infarction and on cardiac function following coronary artery occlusion. Am Heart J 1936;12:285–300.

8. Moss AJ, McDonald J. Unilateral cervicothoracic sympathetic ganglionectomy for the treatment of long QT interval syndrome. N Engl J Med 1971;285:903–4.

9. Fiorelli A, D’Aponte A, Canonico R et al. T2–T3 sympathectomy versus sympathicotomy for essential palmar hyperhidrosis: comparison of effects on cardio-respiratory function†. European Journal of Cardio-Thoracic Surgery 2012;42:454–461.

10. Ibrahim M, Menna C, Andreetti C et al. Bilateral single-port sympathectomy: long-term results and quality of life. Biomed Res Int 2013;2013:348017–348017.

11. Schwartz PJ. Cardiac sympathetic denervation to prevent life-threatening arrhythmias. Nature reviews Cardiology 2014;11:346–53.

12. Zanoni FL, Simas R, da Silva RG et al. Bilateral sympathectomy improves postinfarction left ventricular remodeling and function. J Thorac Cardiovasc Surg 2017;153:855–863.

13. Krishnan A, Suarez-Pierre A, Crawford TC et al. Sympathectomy for Stabilization of Heart Failure Due to Drug-Refractory Ventricular Tachycardia. Ann Thorac Surg 2018;105:e51–e53.

14. Richardson T, Lugo R, Saavedra P et al. Cardiac sympathectomy for the management of ventricular arrhythmias refractory to catheter ablation. Heart Rhythm 2018;15:56–62.

15. Zhou M, Liu Y, Xiong L et al. Cardiac Sympathetic Afferent Denervation Protects Against Ventricular Arrhythmias by Modulating Cardiac Sympathetic Nerve Activity During Acute Myocardial Infarction. Med Sci Monit 2019;25:1984–1993.

16. Aziz H, Nathoo N, Ajlan M et al. Bilateral Cardiac Sympathectomy and Extrapericardial Coil Implantation for the Management of Electrical Storm. JACC Case reports 2021;3:491–495.

17. Ardell JL, Foreman RD, Armour JA, Shivkumar K. Cardiac sympathectomy and spinal cord stimulation attenuate reflex-mediated norepinephrine release during ischemia preventing ventricular fibrillation. JCI Insight 2019;4.

18. Treatment of Ventricular Arrhythmia by Permanent Atrial Pacemaker and Cardiac Sympathectomy. Ann Intern Med 1968;68:591-597.

19. Vaseghi M, Barwad P, Malavassi Corrales FJ et al. Cardiac Sympathetic Denervation for Refractory Ventricular Arrhythmias. J Am Coll Cardiol 2017;69:3070–3080.

20. Meng L, Shivkumar K, Ajijola O. Autonomic Regulation and Ventricular Arrhythmias. Current treatment options in cardiovascular medicine 2018;20:38.

21. Boukens BJD, Dacey M, Meijborg VMF et al. Mechanism of ventricular premature beats elicited by left stellate ganglion stimulation during acute ischaemia of the anterior left ventricle. Cardiovascular research 2021;117:2083–2091.

22. Butrous GS, Gough WB, Restivo M, Yang H, el-Sherif N. Adrenergic effects on reentrant ventricular rhythms in subacute myocardial infarction. Circulation 1992;86:247–54.

23. Yalin K, Liosis S, Palade E et al. Cardiac sympathetic denervation in patients with nonischemic cardiomyopathy and refractory ventricular arrhythmias: a single-center experience. Clinical Research in Cardiology 2020.

24. Tygesen H, Wettervik C, Claes G et al. Long-term effect of endoscopic transthoracic sympathicotomy on heart rate variability and QT dispersion in severe angina pectoris. Int J Cardiol 1999;70:283–92.

25. Assis FR, Sharma A, Shah R et al. Long-Term Outcomes of Bilateral Cardiac Sympathetic Denervation for Refractory Ventricular Tachycardia. JACC Clin Electrophysiol 2021;7:463–470.

26. Dey S, DeMazumder D, Sidor A, Foster DB, O’Rourke B. Mitochondrial ROS Drive Sudden Cardiac Death and Chronic Proteome Remodeling in Heart Failure. Circ Res 2018;123:356–371.

27. Stengl M. Experimental models of spontaneous ventricular arrhythmias and of sudden cardiac death. Physiol Res 2010;59 Suppl 1:S25–31.

28. Lewis GD, Docherty KF, Voors AA et al. Developments in Exercise Capacity Assessment in Heart Failure Clinical Trials and the Rationale for the Design of METEORIC-HF. Circulation: Heart Failure 2022;15:e008970.

29. Allen BJ, Casey TP, Brodsky MA, Luckett CR, Henry WL. Exercise testing in patients with life-threatening ventricular tachyarrhythmias: Results and correlation with clinical and arrhythmia factors. American heart journal 1988;116:997–1002.

30. Yang JC, Wesley RC, Jr, Froelicher VF. Ventricular Tachycardia During Routine Treadmill Testing: Risk and Prognosis. Archives of Internal Medicine 1991;151:349–353.

31. Frolkis JP, Pothier CE, Blackstone EH, Lauer MS. Frequent Ventricular Ectopy after Exercise as a Predictor of Death. New England Journal of Medicine 2003;348:781–790.

32. Dey S, Sidor A, O’Rourke B. Compartment-specific Control of Reactive Oxygen Species Scavenging by Antioxidant Pathway Enzymes. The Journal of biological chemistry 2016;291:11185–97.

33. DeMazumder D, Kass DA, O’Rourke B, Tomaselli GF. Cardiac resynchronization therapy restores sympathovagal balance in the failing heart by differential remodeling of cholinergic signaling. Circ Res 2015;116:1691–9.

34. DeMazumder D, Limpitikul WB, Dorante M et al. Entropy of cardiac repolarization predicts ventricular arrhythmias and mortality in patients receiving an implantable cardioverter-defibrillator for primary prevention of sudden death. Europace 2016;18:1818–1828.

35. Kourie JI. Interaction of reactive oxygen species with ion transport mechanisms. The American journal of physiology 1998;275:C1–24.

36. Zima AV, Blatter LA. Redox regulation of cardiac calcium channels and transporters. Cardiovascular research 2006;71:310–21.

37. Terentyev D, Gyorke I, Belevych AE et al. Redox modification of ryanodine receptors contributes to sarcoplasmic reticulum Ca2+ leak in chronic heart failure. Circ Res 2008;103:1466–72.

38. Liu T, Yang N, Sidor A, O’Rourke B. MCU Overexpression Rescues Inotropy and Reverses Heart Failure by Reducing SR Ca(2+) Leak. Circ Res 2021;128:1191–1204.

39. Ho HT, Stevens SC, Terentyeva R, Carnes CA, Terentyev D, Gyorke S. Arrhythmogenic adverse effects of cardiac glycosides are mediated by redox modification of ryanodine receptors. The Journal of physiology 2011;589:4697–708.

40. Xie A, Song Z, Liu H et al. Mitochondrial Ca(2+) Influx Contributes to Arrhythmic Risk in Nonischemic Cardiomyopathy. J Am Heart Assoc 2018;7.

41. Liu M, Liu H, Dudley SC, Jr. Reactive oxygen species originating from mitochondria regulate the cardiac sodium channel. Circ Res 2010;107:967–74.

42. Liu M, Sanyal S, Gao G et al. Cardiac Na+ current regulation by pyridine nucleotides. Circ Res 2009;105:737–45.

43. Droge W. Free radicals in the physiological control of cell function. Physiol Rev 2002;82:47–95.

44. Song M, Chen Y, Gong G, Murphy E, Rabinovitch PS, Dorn GW, 2nd. Super-suppression of mitochondrial reactive oxygen species signaling impairs compensatory autophagy in primary mitophagic cardiomyopathy. Circ Res 2014;115:348–53.

